# miR126-mediated impaired vascular integrity in Rett syndrome

**DOI:** 10.1101/2024.10.11.617929

**Authors:** Tatsuya Osaki, Zhengpeng Wan, Koji Haratani, Ylliah Jin, Marco Campisi, David A. Barbie, Roger Kamm, Mriganka Sur

## Abstract

Rett syndrome (RTT) is a neurodevelopmental disorder that is caused by mutations in melty-CpG binding protein 2 (MeCP2). MeCP2 is a non-cell type-specific DNA binding protein, and its mutation influences not only neural cells but also non-neural cells in the brain, including vasculature associated with endothelial cells. Vascular integrity is crucial for maintaining brain homeostasis, and its alteration may be linked to the pathology of neurodegenerative disease, but a non-neurogenic effect, especially the relationship between vascular alternation and Rett syndrome pathogenesis, has not been shown. Here, we recapitulate a microvascular network using Rett syndrome patient-derived induced pluripotent stem (iPS) cells that carry MeCP2[R306C] mutation to investigate early developmental vascular impact. To expedite endothelial cell differentiation, doxycycline (DOX)-inducible ETV2 expression vectors were inserted into the AAVS1 locus of Rett syndrome patient-derived iPS cells and its isogenic control by CRISPR/Cas9. With these endothelial cells, we established a disease microvascular network (Rett-dMVNs) and observed higher permeability in the Rett-dMVNs compared to isogenic controls, indicating altered barrier function by MeCP2 mutation. Furthermore, we unveiled that hyperpermeability is involved in the upregulation of miR126-3p in Rett syndrome patient-derived endothelial cells by microRNA profiling and RNAseq, and rescue of miR126-3p level can recover their phenotype. We discover miR126-3p-mediated vascular impairment in Rett syndrome patients and suggest the potential application of these findings for translational medicine.

## Introduction

Rett syndrome (RTT) is a neurodevelopmental disorder that is caused by mutations in the X-linked gene encoding melty-CpG binding protein 2 (MeCP2) ^1,2^. Patient symptoms typically are recognized in early developmental stage between 6-12 months including the loss of hand skills, speech and social engagement, motor abnormalities, and cognitive impairments ^3^. MeCP2 is expressed in various types of cells including neural cells (neurons and astrocytes) and non-neural cells such as endothelial cells and pericytes. Because of pervasiveness of MeCP2 expression and the melty-CpG binding site, its mutation influences not only neuronal health and activity but also non-neural tissue in the brain, including vascular networks associated with endothelial cells ^4,5^. Indeed, RTT patients frequently present with reduced skeletal growth, hypo-perfusion in the area of the midbrain and upper brainstem, and poor circulation ^6,7^, implicating a compromised brain microvasculature in RTT. In addition, vasculogenesis and angiogenesis toward the neural tube are some of the primitive events in neudodevelopment and it occurs before neurogenesis and oligodendrocytegenesis as well as astrocytogenesis ^8^. Vascular formation and differentiation in the brain begins during the early embryonic period and it becomes functional shortly once it is formed ^9–11^ instead astrocytes and myelinated neurons do not appear until soon after birth^12^. Therefore, primitive vascular alternation accompanied by developmental neurotoxicity at a systems biology level widely impacts following neurodevelopmental events and disease progression^13–15^.

Vascular permeability is the capability to control the molecule exchange between vessels, tissues, and organs. In the brain, specifically, the blood-brain barrier (BBB) strictly regulates molecule transportation, including nutrients, waste, and toxins in both influx and efflux directions to maintain homeostasis of the brain in health conditions^16^ and its alternations are linked to a pathological process in neurodegenerative diseases such as Alzheimer’s disease ^17,18^ and Parkinson’s disease ^19^ ^20^. Interestingly, it has been reported this BBB breakdown is present prior to Aβ plague accumulation in Alzheimer’s disease patients ^21^. Therefore, BBB breakdown may trigger the alternation of neuronal circumstances, resulting in induced neurodegeneration, mitochondrial dysfunction as well as aggregation of Aβ, but no pathological role of vascular alternation in RTT has been shown so far.

To date, there has been only limited evidence indicating impaired BBB permeability in both Rett syndrome individuals and MeCP2-null mouse models, but existing evidence is growing and accumulating. Recent studies showed that MeCP2-null mouse (Mecp2-/y, Mecp2^tm1.1Bird^) exhibited alternation of BBB integrity along with the decreased expression of tight junctions such as OCLN and CLDN5 in cortex ^22^. It might be impacted by increased matrix metallopeptidase 9 (MMP9) expression by inducing the degradation of basal lamina. Vessels from female MeCP2^+/−^ mice show a reduced endothelium-dependent relaxation due to a reduced Nitric Oxide (NO) availability secondary to an increased reactive oxygen species generation ^23^. Additionally, RNA sequencing and proteomics approaches reveal morphology alteration of the blood vessel in Mecp2^tm^^1.1Jae/+^ mice and increase the protein level related to the blood-brain barrier ^24^. Therefore, altered brain vascular homeostasis and subsequent BBB breakdown may be induced in MeCP2 deficient mice (MeCP2 null mice). However, the specific role of MeCP2 mutation in human development with respect to vascular permeability and association with neurodevelopmental alternation accompanying neurovascular interaction remains unknown due to the lack of a disease model recapitulating this complicated pathological process.

Here, we engineer a 3D microvascular network (MVN) model in microfluidic devices using Rett syndrome patient-derived induced pluripotent stem (iPS), which carries MeCP2[R306C] mutation and investigate endothelial specific molecular pathway by MeCP2 mutation undelaying alternation of vascular permeability associated with vascular-specific microRNA (miR-126). MeCP2[R306C] mutation on the transcription repression domain (TRD) is one of the common SNP mutations (9-10%) that does not influence affinity to the methyl CpG binding domain, but it disrupts the interaction with the NCoR corepressor complex ^25^. We first modified the iPS cell from the Rett syndrome patient by knocking in doxycycline (DOX)-inducible ETV2 gene cassette by CRISPR/ Cas9, which is now one of the promised ways to directly differentiate endothelial cells rapidly ^26,27^. iEC were then embedded in a fibrin gel in a microfluidic device to form 3D vascular networks, which were subsequently used for permeability tests. Impaired vascular permeability was observed in patient-derived vasculature with Rett syndrome. Subsequent transcriptomic analysis elucidated that these impartments are associated with the signaling pathway of miR-126, along with the decreasing tight junction expression mediated by these impairments. Moreover, downregulating miRNA-126-3p by expressing antisense miRNA-126-3p partially restored endothelial barrier function and upregulated tight junctions, inhibiting the Ang2/Tie2 signaling pathway. These results provide clear evidence that miR-126-3p is a direct target of MeCP2 and a mediator of vascular impairment caused by MeCP2[R306C] mutations. It also identifies miR-126-3p as a promising therapeutic target for restoring vascular impairment. This study holds promise for potential translational applications aimed at rescuing vascular alterations in the early developmental stages of Rett syndrome, as well as in other neurodegenerative and developmental disorders.

## RESULTS

### ETV2 expressing iPS cells derived from Rett syndrome patient

To recapitulate the vascular formation process in early development in Rett syndrome and assess vascular permeability, we established the stable iPS cells line from Rett syndrome patient, which has Doxycycline (DOX)-inducible ETV2 expression to obtain iPS-induced endothelial cells (iEC, **Fig. 1a**). The all-in-one plasmid that expresses ETV2 under rtTA and Tet3G under CAG bidirectional promoter with mCherry and AAVS1 targeting sgRNA with SpCas9 were constructed from pUCM-AAVS1-TO-hNGN2 plasmid and then it delivered to Rett syndrome patient-derived iPS cells which carry MeCP2 [R306C] mutation (916C>T) and isogenic control iPS cells by electroporation (**Fig. 1b**). SNP mutation in iPS cells were confirmed by Sanger sequencing followed PCR (**Supplemental fig. 1a**). Genome edited iPS cells were further sorted by cell sorted with mCherry positive cells and puromycin selection. Although mCherry-positive cells are less than 5% before sorting, we obtained a 100% positive population by cell sorting. To test the response of ETV2 expression in the presence of DOX, we treated DOX in iPS cells for 12, 24, 48, and 72 hours and then extracted the protein. Western blotting results revealed that ETV2 in genome-edited cells started to be expressed in 12 hours and reached fully expressed in 48 hours, although very low leak expression of ETV2 was observed in the absence of DOX in both iPS cell line due to bi-directional promoters and 3^rd^ generation Tet-on system (**Supplemental fig. 1b**). Pluripotency of iPS cells after genome editing were further characterized by immunostaining with Oct3/4, Nanog, SSEA1, TRA-1-60, and SSEA4, which suggested that no unexpressed differentiations were observed (**Supplemental fig. 1c**).

**Figure 1.**
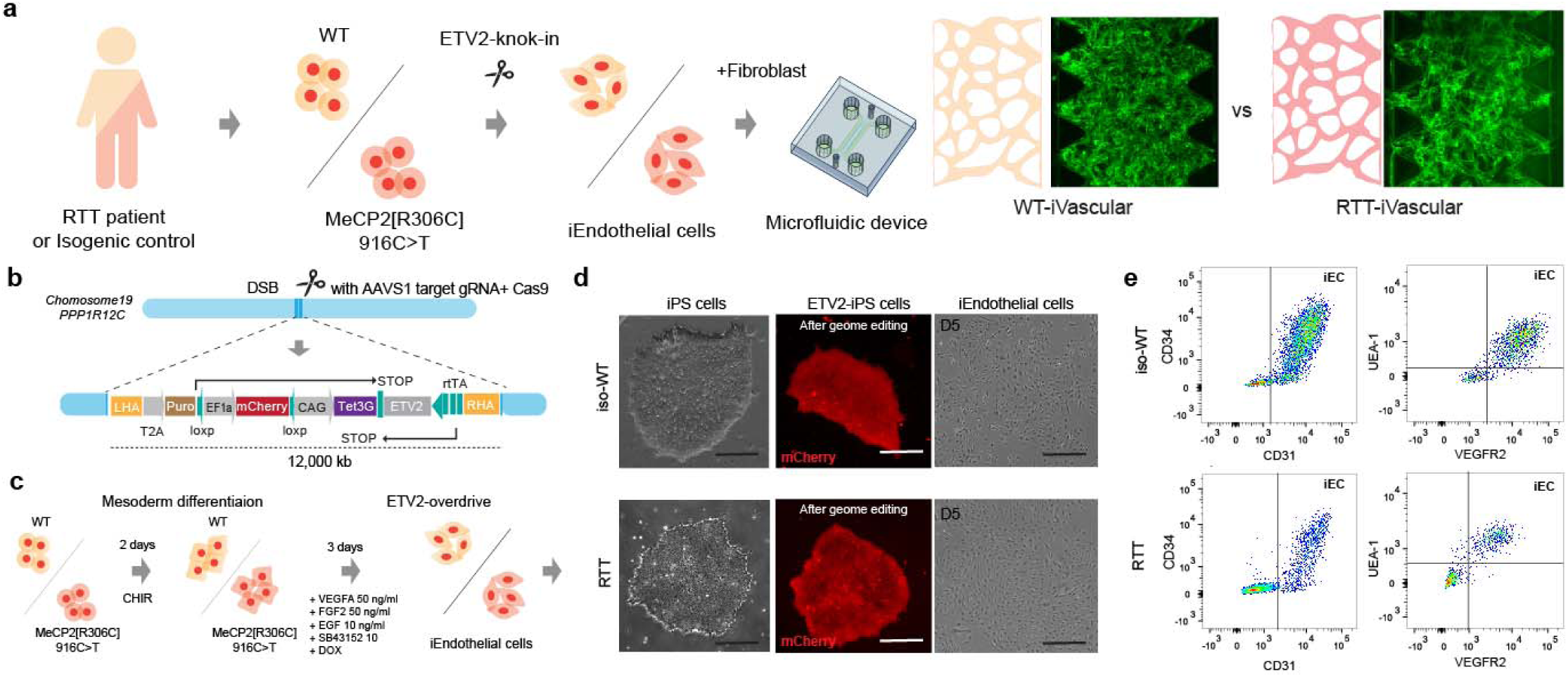
|CRISPR-ETV2 knock-in line in Rett syndrome patient-derived iPS cells. (**a**) Schematic illustration of engineering Rett syndrome’s microvascular network in the microfluidic device. Rett syndrome patient-derived iPS cells which carry MECP2[R306C] mutations and isogenic control were modified by knocking in ETV2-gene cassette by CRISPR/Cas9 to accelerate the endothelial differentiation. Then, induced endothelial cells (iEC) and fibroblast were introduced to the microfluidic device to form microvascular networks. (**b, c, d**) ETV2 under Tet3G promoter cassette was inserted to AAVS1 safe harbor locus and cells were sorted by mCherry fluorescent marker by FACS to obtain stable iPS cell line (**d**). iECs were obtained in 5 days from iPS cells in the presence of DOX. (**d**) Single cloned iPS cells (Rett syndrome patient-derived and control) with ETV2 gene cassettes (mCherry) and representative images of morphology of endothelial cells after 5 days from the differentiation. (**e**) Endothelial differentiation was confirmed by flow cytometery according to CD31, CD34, VEGFR2, and UEA-1 expression.

### Differentiation and characterization of induced endothelial cells (iEC)

The ETV2-knock in iPS cells were then, treated with GSK3 inhibitor/Wnt activator (CHIR99021) for two days for mesoderm differentiation, and then treated with VEGF, EGF, SB43152, and Doxycycline for 3 days to differentiate into induced iEC (**Fig. 1c**). Flow cytometry showed that iEC were expressed with CD31, CD34, UEA1, and VEGFR2 that are typical endothelial markers in both RTT patient-derived endothelial cells and its isogenic control. iEC were characterized by immunostaining with ZO-1 and quantified by area fraction of ZO-1 to clarify tight junction. In RTT-iEC, ZO-1 expressions decreased and did not localize at the boundary between cells compared to isogenic control (**Fig. 2A**). Area fraction of ZO-1 in RTT-iEC was also lower than in control (**Fig. 2b**). RT-PCR results suggested that gene expressions involved in tight junction, such as ZO-1, OCLDN, CLDN5, were down-regulated (**Fig. 2c and d**). RTT-iECs also downregulated the gene expression of transporters, including MCT1, PGP1, LRP1, and GLUT1, compared to WT-iECs and HUVECs (human umbilical vein endothelial cell, **Fig. 2c**). We also measured trans-endothelial electrical resistance (TEER) to assess barrier function in a 2D monolayer of endothelial cells. As a result, TEER of RTT-iEC were lower than WT-iEC and HUVEC on 21 days of culture (**Fig. 2e**). RNAseq revealed that down-regulated (1162, p<0.05) and up-regulated genes (1163, p<0.05) and global effect of MeCP2[R306C] mutation in RTT-iEC (**Fig. 2f and g**). We also found that ZO-1 expression was downregulated in RTT-iEC (**Fig. 2f**). According to GO enrichment analysis, molecular pathways with respect to biological processes related to “Regulation of vascular development” and “Protein localization to cell-cell junction” were down-regulated in RTT-iEC (**Fig. 2h**).

**Figure 2.**
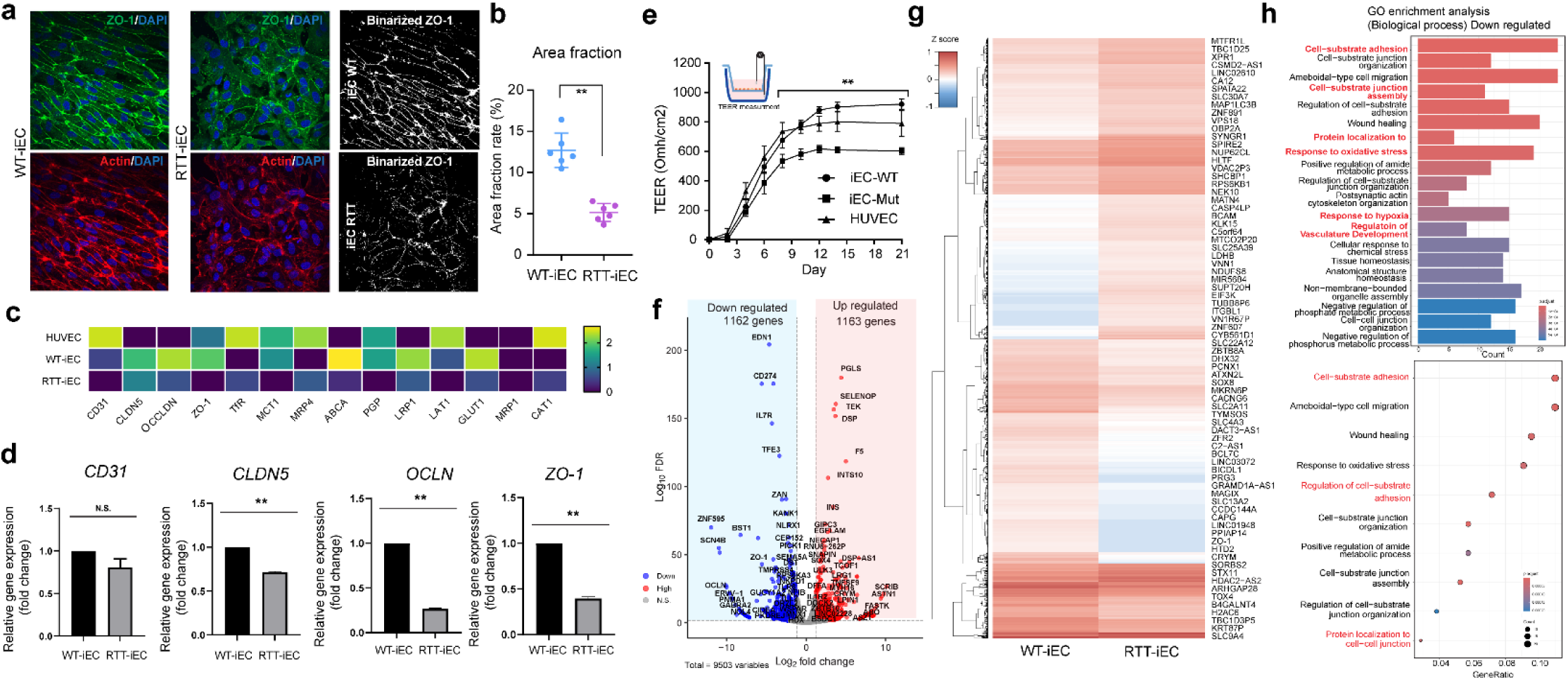
|Characterization of iEndothelial cells in 2D culture. (**a**) Immunostaining of ZO-1 (green) and F-actin (red) to quantify tight junctions’ property (**b**) Quantification of ZO-1 area fraction rate. iEC-Mut displayed lower area fractions than iEC-WT. (**c**) Gene expression showed (**d**) iEC-Mut exhibited lower gene expressions related to tight junctions (CLDN5, OCLN, and ZO-1) measured by PCR. n = 3. (**e**) Measurement of TEER indicating endothelial barrier functions. (**f, g**) Differential gene expression by RNAseq. n = 2. (**h**) GO enrichment analysis. Student’s t-test, **, *p*<0.01, * *p*<0.05.

For the first time, we established a protocol to differentiate RTT-ECs and fully characterize them by RNAseq. To further quantify the endothelial characterization, imaging-enhanced flow cytometry was performed (**Fig. 3a**). It allows us to validate morphological character on top of cell surface marker staining (ZO-1). The flowed cells were gated to identify single cells judging from acquired imaging and combined FSC and SSC values, then the cell cycle was evaluated with DAPI staining. RTT-iEC showed clear S phase arrest from G0/G1 phase, although WT-iEC represented a typical cell cycle phase (**Fig. 3b, c**). Perimeter, pseudo-diameter of cells, entropy, and circularity were then calculated from images, and they were clustered and visualized as a UMAP plot (**Fig. 3d, e**). We classified 6 identical clusters based on these 4 parameters and overlaid other information (cell cycle tag, mutation of MeCP2, and ZO-1 expression from immune flow chemistry, **Fig. 3f**). We classified ZO-1 highly expressed clusters (Cluster 3 and 4) and typical endothelial cell clusters (Cluster 2 and 5), and other cell types or dead cells (Cluster 1 and 6, **Fig. 3g**). Population of WT cell in cluster 3 which is high ZO-1 expressing clusters was higher than RTT-iEC and cluster 5 population which is typical endothelial cell clusters was lower (**Fig. 3h, i, and j**). These results showed that RTT-iEC could be differentiated into endothelial cells that express lower ZO-1 compared to WT-iEC and suggested that MeCP2 R306C mutation itself negatively impacted the integrity of tight junction in differentiated endothelial cells.

**Figure 3.**
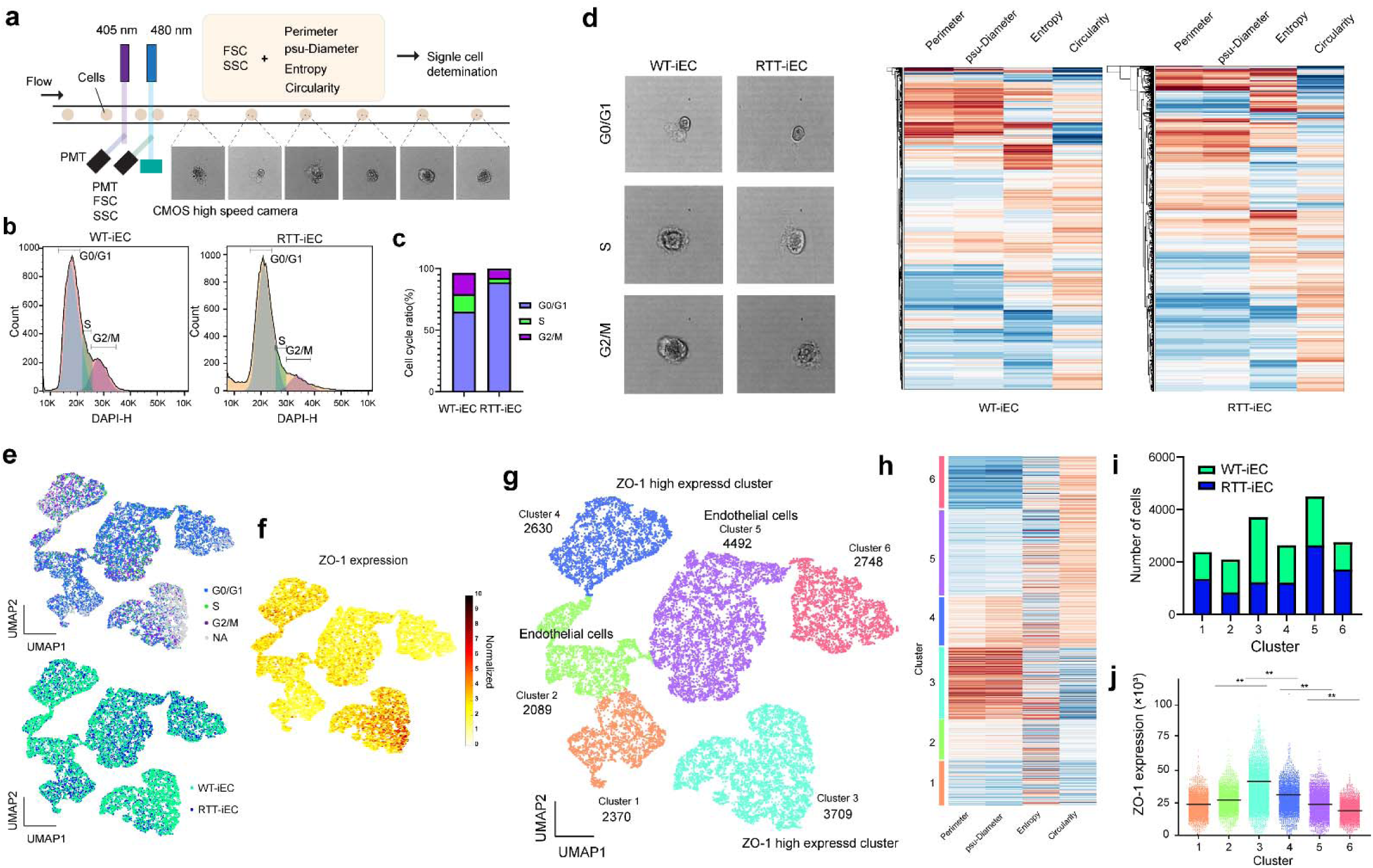
Imaging enhanced flow cytometry revealed heterogeneity of mutant endothelial cells. **(a)** Schematic illustration of imaging enhanced flow cytometry. (**b, c**) Single cells gating and CD31 positive cells were determined. (**d**) Cell cycle analysis with DNA staining. (**e**) iEC-RTT had higher population of G0/G1 cycle. (**f**) Representative image of iEC-WT and iEC-Mut according to different cell cycles. Perimeter, psudo-diameter, entropy, and circularity were quantified by images. (**g**) UMAP plot showed the diverse population of endothelial cells. (**h, i**) Feature plot of ZO-1 indicates the unique clusters (cluster 3), which express higher ZO-1. We determined endothelial cell clusters (cluster 2, 4, 5) and ZO-1 high expressed cluster (cluster 3). Clusters were determined by four morphological features (perimeter, psudo-diameter, entropy, and circularity). (**k**) Quantification of number of cells in each cluster. Number of cells in cluster 3 of iEC-WT is higher than iEC-RTT, which expre4ss higher ZO-1.

Of note, condition medium was collected from iEC and added to the iPS-derived neuron (NGN2-driven) and then we performed calcium imaging to analyze the differential impact of secreted protein from endothelial cells to neuron cells (**Supplemental fig. 2a, b**). Treatment with condition medium from WT-iEC and RTT-iEC did not affect the neurite length and number of cells (**Supplemental fig. 2c**), however, condition medium from RTT-iEC negatively impacted the neuronal activity in healthy neurons (**Supplemental fig. 1d, e, and f**). This result suggested altered neurovascular coupling mediated by a paracrine signal from endothelial cells to neurons.

### Impairment of vascular permeability in Rett syndrome patient derived 3D microvascular networks

According to the characterization of 2D monolayer and gene expression analysis, RTT-iEC exhibited the altered gene expression of tight junction. Although 2D TEER measurements is the promised method to estimate the paracellular barrier function of endothelial cells, there is no specificity of molecules and other factors (e.g., the thickness of layer, morphology, and density) that impact the variation of the outcome. To classify true vascular permeability, we engineered a 3D vascular network in a microfluidic device with WT-iEC or RTT-iEC. The cells were embedded in fibrin gel with healthy human lung fibroblasts as supporting cells and introduced into the microdevice’s center channel. After gelation for 10 min, a culture medium was introduced into the side channel and cultured for 5 days, resulting in the formation of the microvascular network in the microfluidic device (**Fig. 4a**). In both WT-iEC and RTT-iEC could form 3D perusable microvasculature in 5 days (WT-iVascular and RTT-iVascular). Notably, RTT-iVascular formed slightly smaller diameter of microvasculature than WT-iVascular **Fig. 4b**. Texas-red conjugated 70kDa dextran perfusion further visualize the perfusability and circularity of microvasculature, whereas no significant difference was observed in the roundness of each vasculature (**Fig. 4c**). To measure permeability of endothelial barrier in 3D microvasculature, vasculature with 70 kDa of dextran were imaged subsequently 10 min later right after dextran injection (**Fig. 4d**). RTT-iVascular showed higher permeability compared to WT-iVascular (**Fig. 4e**). Immunostaining with ZO-1 in 3D microvasculature also demonstrated that WT-iVascular formed more packed vasculature along with clear boundary and higher intensity of ZO-1 than RTT-iVascular (**Fig. 4f**). In addition, we stained MVNs with collagen IV to characterize extracelluer matrix and observedno significant differences (**Fig. 4g**). We collected the culture medium from the 3D device and quantified MMP and TIMP level, because these secreted proteins play an important role to vasculogenic and remodeling of vasculature in developmental process and MeCP2 null mutation influenced the secretion level of MMP9 ^22^. Unlike previous studies using mouse model, no significant difference in MMP9 level was observed between WT-iVascular and RTT-iVascular, however, we observed clear differences of MMP8, TIMP1, and TIMP4 (**Fig. 4h**). MMP8 is also one of the crucial collagen cleaving enzymes, and TIMP1 is also known as the natural inhibitor of the MMPs ^28^. Impaired MMP8 and TIMP1 activity in RTT-iVascular should be one of the triggers for decreased barrier function measured by permeability assay in 3D conditions. Next, we also engineered 3D blood-brain barrier (BBB) in a microfluidic device using WT-iEC or RTT-iEC, astrocyte and pericyte (**Supplemental fig. 3a**). All three types of cells were embedded in fibrin gel and introduced into the center channel of microdevice, resulting in the formation of BBB formation (**Supplemental fig. 3b**). Same as vasculature, BBB which express mutant MeCP2 in endothelial cells exhibited higher permeability (**Supplemental fig. 3c**).

**Figure 4.**
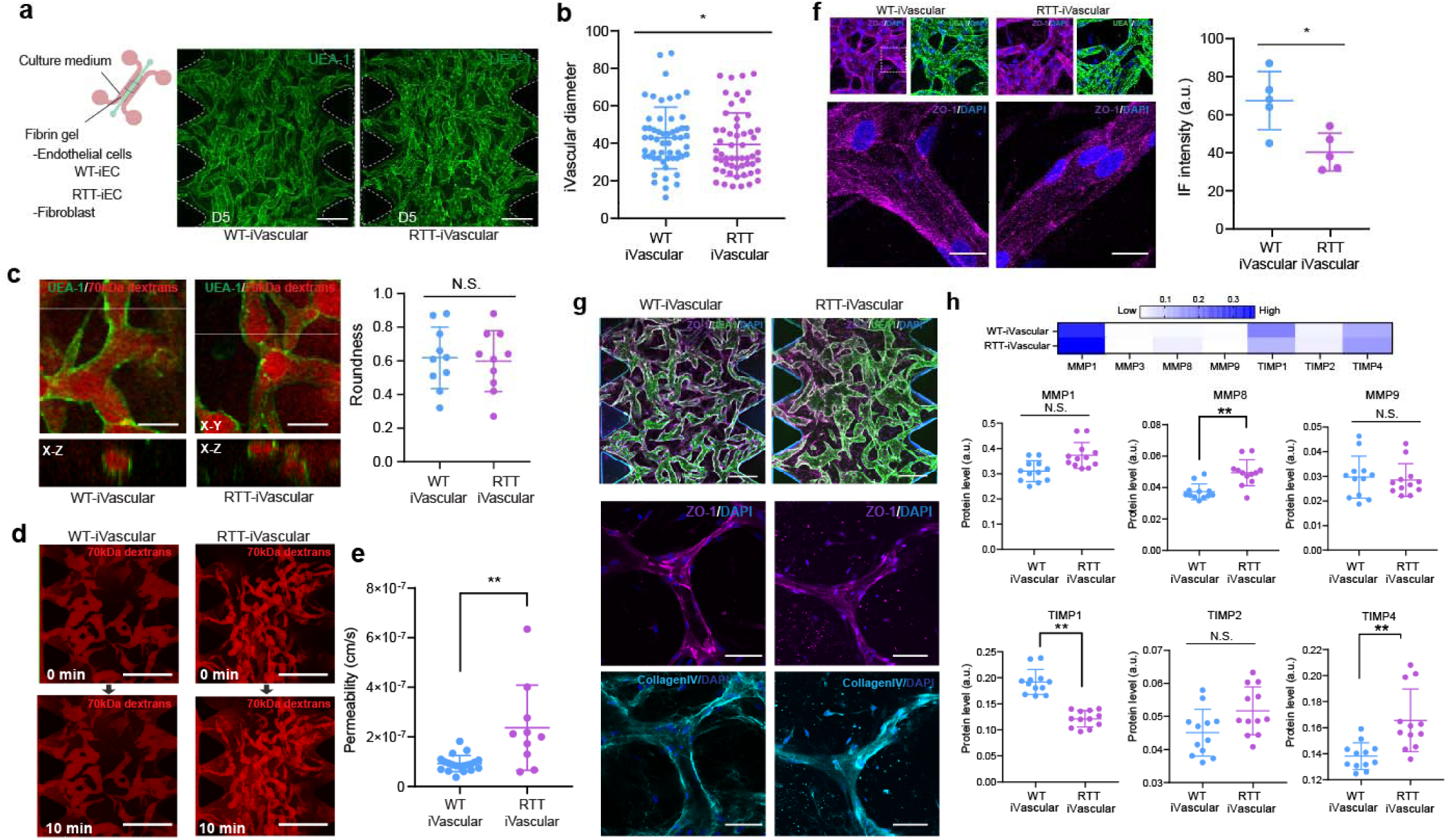
Rett syndrome patient-derived microvascular network exhibited higher permeability along with altered tight junction expression. (**a**) WT-iEC and RTT-iEC were embedded in fibrin gel with fibroblast and introduced into microfluidic device to form microvascular networks. Regardless of MeCP2 mutation, microvascular networks were formed with in 5 days. (**b**) RTT-iVascular exhibited slightly but significantly smaller diameter of vasculature. (**c**) There is no significant difference of roundness of vasculature. (**d**) Permeability of endothelial layer were quantified by perfusion of 70 kDa of Texas-red conjugated dextran solution. (**e**) RTT-iVascular exhibited higher permeability indicating lower barrier function of endothelial layer. (**f**) Immunostaining of ZO-1 in 3D microvascular networks. ZO-1 intensity in RTT-iVascualr was lower than WT-iVascular. (**g**) Both WT-iVascular and RTT-iVascular produced collagenIV, but no significant difference. (**h**) MMP and TIMP1 levels were quantified from condition medium in 3D microvasculature. RTT-iVacular secreted higher MMP8 and TIMP4 and lower TIMP1, but no significant difference in MMP1 and MMP9. Student’s t-test, **, *p*<0.01, * *p*<0.05.

### Pathological mechanism associated with endothelial-specific miR-126 dysregulation

microRNAs are small noncoding RNAs that function as post-transcriptional regulators of gene expression. They play a vital role in numerous biological processes, including proliferation, differentiation, and development in posttranslational level ^29^. The involvement of MeCP2 in the regulation of miRNA expression was first reported in 2010 ^30,31^. Indeed, miR-199a has been identified to control the functions of neurons mediated by mTOR signaling pathway ^32^. However, whether this MeCP2/miRNA pathway is implicated in other aspects of the vascular integrity in Rett syndrome remains unexplored. To validate how MeCP2 mutation results in decreased tight junction protein, we focused on microRNA expression, especially endothelial cell-specific microRNA, miR-126-3p, in 2D cultured endothelial cells (**Supplemental fig. 4a**). Total microRNA was isolated from WT-iEC and RTT-iEC in 2D culture, and then microRNA expressions were quantified by RT-PCR with appropriate internal control (**Supplemental fig. 4b and c**). As a result, miR-126-3p expression in RTT-iEC is significantly higher than WT-iEC (**Fig. 5a**). miR-126-3p is only expressed in endothelial cells and acts upon various transcripts to control angiogenesis ^33^. Several target genes were displayed by MirTarget prediction algorithm ^34^ (**Fig. 5b**). *VEGF*-A, *SPRED1*, and *PIK3R2* play an especially important role in controlling vascular integrity. RT-PCR results showed that *EGFL7*, *IGFBP2*, *BEGFA*, *ADAM9*, *SPRED1*, and *PIK3R2* were downregulated, but not *HIF1A* and *NFKBIA* in RTT-iEC compared to WT-iEC (**Fig. 5c**).

**Figure 5.**
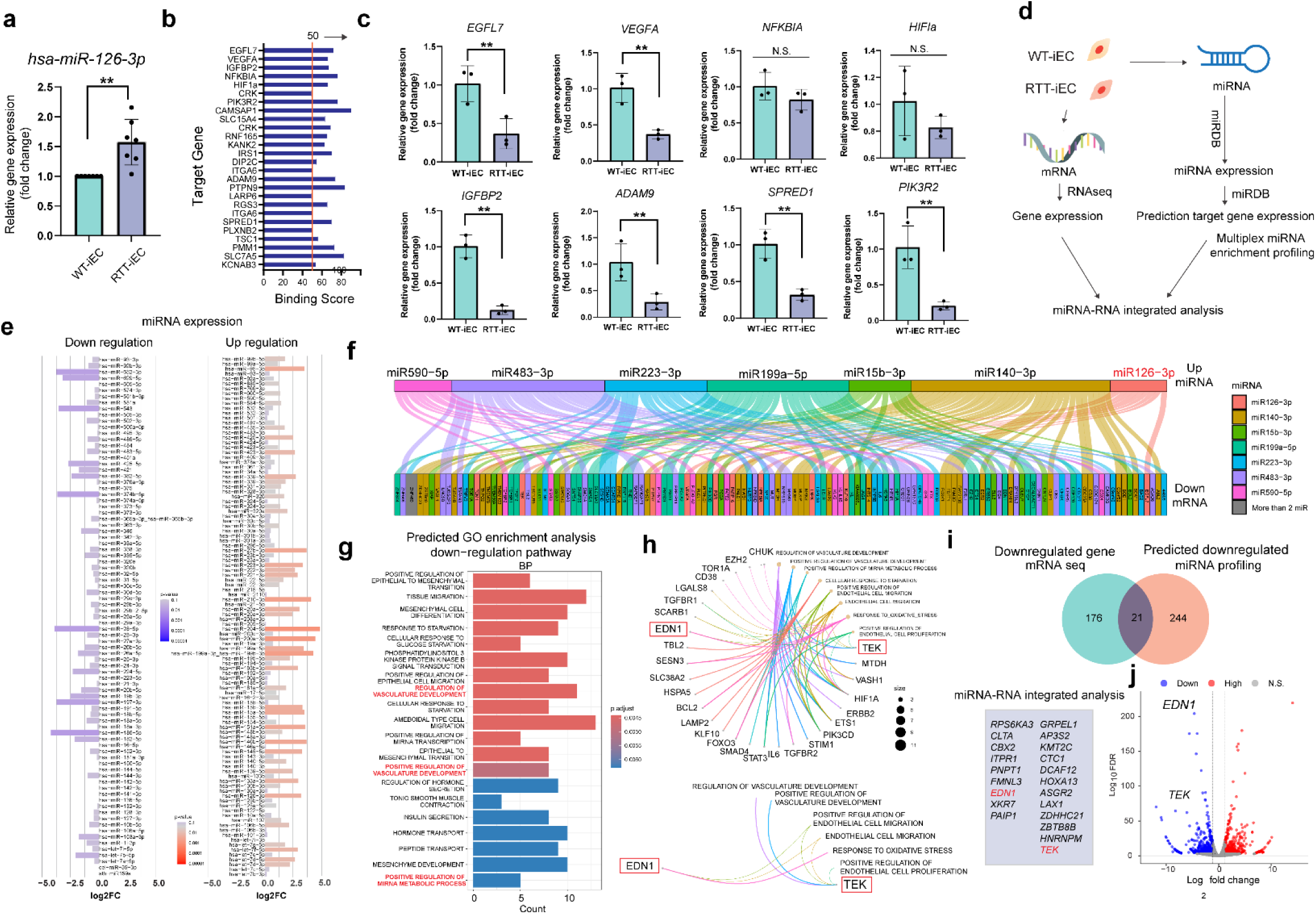
Upregulation of microRNA-126 in RTT-iEC negatively impacted vascular development and permeability. **(a)** miR-126-3p was highly expressed in RTT-iEC compared to isogenic control endothelial cells. (**b**) miRDB predict the target gene of miR126-3p. (**c**) Down regulation of gene expressions, which are targeted by miR126-3p. (**d**) miRNA-integrated analaysis was performed using RNAseqecing data and predicted gene expression measured by PCR based miR expression analysis along with miRDB prediction. (**e**) Downregualation and upregulation of miRNA of RTT-iEC over WT-iEC. (**f**) 7-upregulated microRNA were determed by the miRNA expression analsysis (**e**) and endothelial cell related. (**g**) GO endrichment analysis (biological process) from predicted downregulated gene set determined by miRDB and miRNA expression analsysis (**e**). Significant down regulated pathway (Regulation of vascular development) were determined. (**h**) Cnet plot suggested that gene involved in these signaling pathway including END1 and TEK1. (**i**) miRNA-RNA integrated analysis revelared 21 overlap downregulated genes including EDN1 and TEK1, which are also consinted with RNAseq analysis (**j**), which potential involved in miR126-3p-mediated vascular alternation.

To expand the effect of miRNA expression in mutant endothelial cells, we performed miRNA-mRNA multi-modal analysis to estimate the influence of miRNA expression and target mRNA gene expression (**Fig. 5d**). The differential miRNA expression in RTT-iEC was shown in **Fig. 5e**. We further pick up the reasonable differential miRNA according to p-value (p <0.01) and fold change (more than 2.5 times). miRNAs including has-miR-126-3p, hsa-miR-140-3p, hsa-miR-15b, hsa-miR199a-5p, hsa-miR-223-5p, hsa-miR-483-3p, and has-miR-590-5p were upregulated and hsa-miR-29a-3p, hsa-miR-29c-3p, hsa-miR-363-3p, and hsa-miR-652-3p were found as downregulated miRNAs (**Fig. 5e**). Alluvial plot showed that target genes of differential miRNA according to miRDB target gene list and performed GO enrichment analysis with respect to biological process based on upregulated and down-regulated genes (**Fig. 5f** and **supplemental fig. 4d**). We discovered one overlapped upregulated (Cell cycle G2M phase translation, **supplemental fig. 4e and f**) and three overlapped downregulated molecular pathway (“(positive) regulation of vascular development” and regulation of miRNA metabolism process) from the miRNA-mRNA profiling (**Fig. 5g**). According to Cnetplot as well as overlap genes in miRNA-RNA profiling (**Fig. 5h** and **i**), the downregulation of Angiopoietin-1 receptor (TEK) and endothelin-1 (EDN1) in RTT-iEC were found in both dowregulated gene expression measured by RNAseq and predicted gene expression from miRNA expression and miRDB prediction (**Fig. 5j**). Angiopoetin-1/Tie2(TEK) signaling pathway play a important role for vascular stabilization via phospholation of PI3K/AKT and forkhead box protein (*FOXO1)*, *RhoA*, VE-PTP signaling pathway^35,36^. Endothelin-1 is a vasoconstrictor peptide, which regulates vascular dilation communicating with smooth muscle cells and cell migration of endothelial cells. These mechanobiological sensors involved in both EDN-1 and Ang1/2/Tie2 signaling regulate vascular homeostasis, and its dysregulation leads to destabilization against external stimuli such as shear stress. Therefore, these two genes might play an important role in the dysfunction of barrier function in mutant RTT-iVascular mediated by altered miRNA expression.

Lastly, to confirm miR-126-3p-mediated dysregulation of vascular permeability, antisense of miR126-3p was transfected to RTT-iEC by lentivirus and formed vasculature in a microfluidic device in the same manner, followed by permeability measurement (**Fig. 6a**). Immunostaining of ZO-1 also that miRNA-126-3p knockdown led to the increase in the ZO-1 expression in the 3D microvascular networks (**Fig. 6b**). In addition, conditional knockdown of miRNA-126-3p could partially restore endothelial barrier function characterized by permeability measurement with 70kDa dextrans (**Fig. 6c, d**). To prove how this restoration event happened, we performed RT-PCR focusing on the gene involved in TEK and EDN-1 signaling pathway(**Fig. 6e**). As shown earlier, we confirmed the downregulation of *TEK* and *EDN-1* in RTT-iEC along with the upregulation of *FOXO1* and *ANGPT2* and the regulation of *EGFL7* and *CLDN5*. Knockdown of miR-126-3p can restore the expression level of *TEK*, *EDN-1*, *FOXO1*, *ANGPT2*, and *CLDN5* (**Fig. 6e**). activation of *FOXO1* and *PI3K* works as a positive feedback regulator of Ang-2/Tie2 signaling pathway^35,36^, which leads to junction instability and less tolerance against shear stress along with RhoA and CLDN5 down-regulation (**Fig. 6f** and **g**). EDN-1 is not a direct regulator to modulate the tight junction property, however, endothelin-1, a vasoconstrictor peptide, plays a critical role in adjusting blood pressure with smooth muscle cells and its alternation, is consistent with previous literature^6,7^ causes dysregulation of blood circulation and its restoration should be important the homeostasis of vascular integrity. Therefore, we conclude that the MeCP2[R306C] mutation impacts miR-126-mediated dysfunction of vascular integrity along with TEK and EDN-1 downregulation in the early development process in Rett syndrome, and its restoration would become one of the therapeutic strategies.

**Figure 6.**
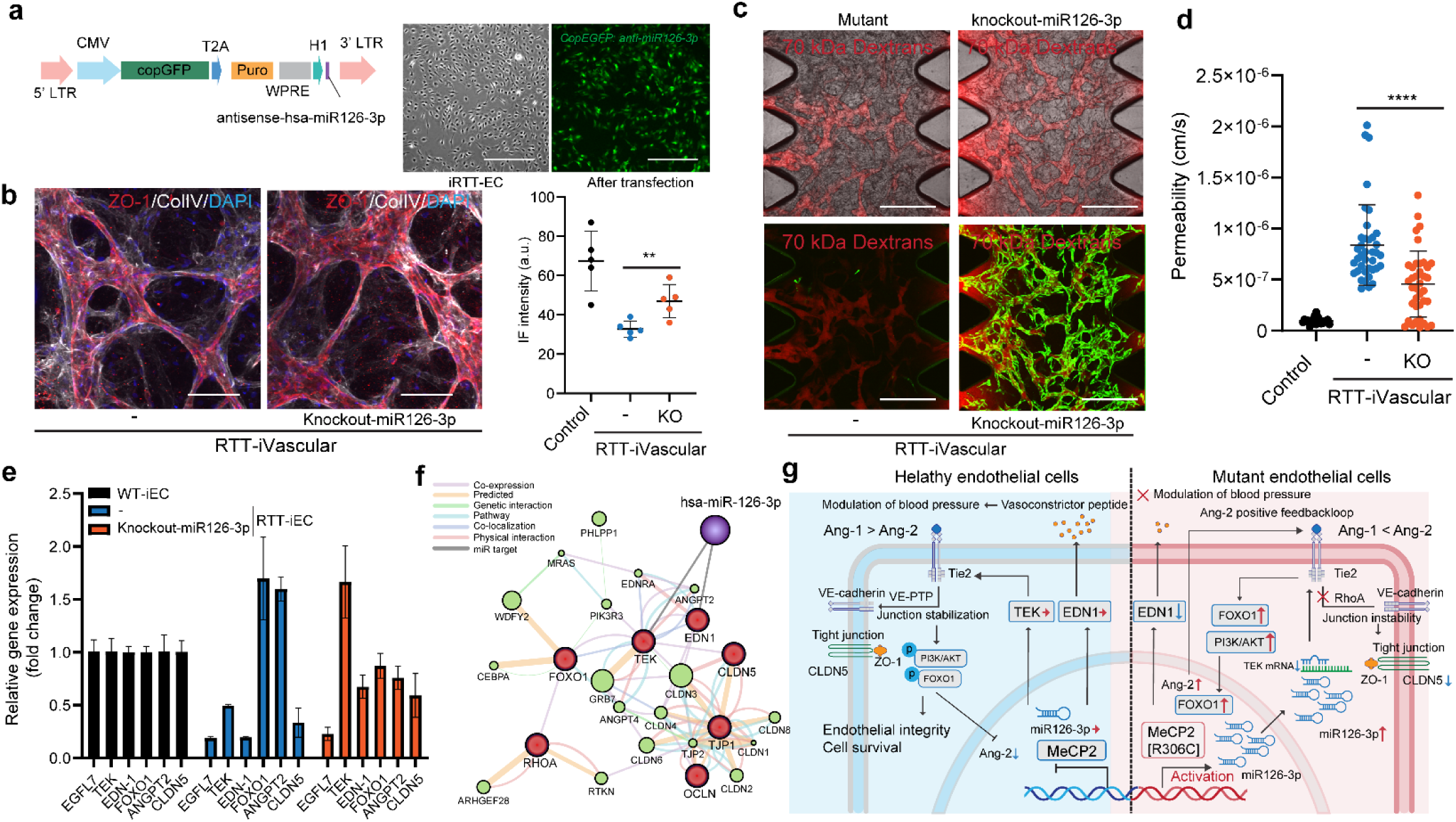
Knockdown of miRNA-126 rescues the phenotype and endothelial barrier functions in RTT-iEC. **(a)** Lentivirus vector to deliver antisense-miR126-3p and EGFP to down-regulate miR126 expressions. (**b**) After endothelial differentiation, the cells were transfected with lentivirus, and EGFP-positive cells were sorted. n =5. (**c, d**) Representative images (c) and pearmeability analysis (d) of RTT-iVascular MVNs with or without expressing miR126-3p knockout constructs (green) perfused with 70 kDa Dextran (red). n =30. (**e**) RT-PCR revealed that EGFL7, TEK, EDN-1, and CLDN5 were downregulated in RTT-iEC, and knockout of miR126-3p recovered TEK and EDN-1 expression along with FOXO1 and CLDN5 rescuing. n =3. (**f**) Gene function prediction involved in Ang/Tie2 signaling and tight junction stabilization based on GeneMANIA algorithm. (**g**) Schematic illustration of altered gene pathway in RTT-iEC. Student’s t-test, ****, *p*<0.0001.

## DISCUSSION

This paper first describes the miRNA-126 mediated vascular alternation in Rett syndrome microvascular networks. We established ETV2-expressing iPS cells that express MeCP2[R306C] mutations and native MeCP2 by CRISPR/Cas9, which allowed us to obtain endothelial cells rapidly (**Fig. 1**), then engineered microvascular networks in a microfluidic device (**Fig. 3**). Location controlled knocking in gene cassette which has Tet-ETV2 into AAVS1 locus realize stable and reproducible endothelial differentiation over lentivirus based ETV2 overexpression^37^ and transfection with mRNA ^38^, allowing them to minimize line-dependent (donor-dependant) variations. Although the procedure of fabricating an MVN has been shown earlier by our group ^39–42^ and promised way, this is the first *in vitro* model that recapitulates early developmental vascular alternation in Rett syndrome, specifically focusing on endothelial barrier function regulating the permeability of molecules by using patient-derived iPS cells. In both 2D and 3D characterization of iEC, tight junction gene expression (ZO-1) was downregulated in endothelial cells and vasculature from Rett syndrome patients (**Fig. 2, 3, and 4**). One of the most important highlights of this study is that we confirmed RTT-iVascular, which is fabricated from Rett syndrome patient-derived iPS cells, displayed higher permeability, indicating altered endothelial barrier functions (**Fig. 4e**). This discovery is consistent with previous literature shown using MeCP2-null mouse (Mecp2-/y, Mecp2tm1.1Bird)^22^ ^23^. Although the MeCP2-null mouse model causes the increase of matrix metallopeptidase 9 (MMP9), and this should be involved in BBB disruption, we did not observe MMP1 and MMP9 dysregulation (**Fig. 4h**), which are critical proteases for vascular turnover with collagen. This might be explained by the structural difference of the non-catalyst domain of MMP2 and MMP9, which is important for substrate binding and interactions between humans and mice ^43–45^, and this suggested the advantage of the human *in vitro* model compared to the mouse model.

Transcriptome analysis revealed that MeCP2[R306C] mutation in endothelial cells itself downregulated as “Regulation of vascular development” and “Protein localization to cell-cell junction” in RTT-iEC (**Fig. 2f, h**). This result advocated that MeCP2 plays an important role in vascular development and homeostasis of vascular integrity, and MeCP2[R306C] mutations negatively impact it. To clarify how MeCP2[R306C] mutations influenced endothelial barrier function along with losing tight junction potential, we focused on endothelial-specific miRNA-126-3p. This miRNA is critical for endothelial differentiation and development ^46,47^ and is also involved in vascular disease ^48^. We discovered miRNA-126-3p is highly expressed in RTT-iEC compared to WT-iEC and target gene expressions of miRNA-126-3p were downregulated in contrast, which contains *EGFL7*, *IGFBP2, BEGFA, ADAM9, SPRED1*, and *PIK3R2 (***Fig.5f**). Integrated analysis of miRNA profiling and RNAseq enforced the evidence showing miRNA-126-3p mediated dysregulation of vascular integrity in RTT-iEC (**Fig. 5f, g**) along with down-regulation of TEK and EDN-1, which are important genes for junction stabilization and cell survival, and regulation of vasoconstriction (**Fig. 5h, i, j and Fig. 6f, g**). Downregulation of TEK further caused the upregulation of FOXO1 and PI3K signaling pathways to activate the Ang2/Tie2 positive feedback loop, leading to vascular instability and decreased vascular integrity (F**ig. 6e**). EDN-1, producing endothelin, is an important driver of vasoconstriction and regulating the blood pressure. Downregulation of EDN-1 in RTT-iEC can explain the poor circulation of blood in RTT patients, which has been shown in previous literature ^6,7^, along with eNOS downregulation^23^. Moreover, the knockdown of miR-126-3p in RTT-iEC could rescue the phenotype of MVNs along with the restoration of TEK and EDN-1 gene expression (**Fig. 6e**). Taken together, these results show clear evidence that miR-126-3p is one of the direct targets of MeCP2 and mediator of vascular impairment by MeCP2[R306C] mutations and one of the good therapeutic target to restore vascular function.

In the transcription process of microRNA, MeCP2 directly interacts with RNA polymerase II to modulate their transcription, and the mutation of MeCP2 disturbs downstream processing ^49,50^. In addition, it has been reported that the interaction between Drosha and DGCR8 along with MeCP2 plays an important role in the processing of miR-199a during the process precursor miRNA (pri-miRNA), and a lack of MeCP2 leads to negatively regulate miR-199 level in neurons, resulting in altered axon development and synapse formation ^32,51^ and neural stem cell fate specification ^52^. Unlike neurons, the role of microRNA in endothelial cells is not known in the context of vascular development. We first observed the microRNA-mediated role of MeCP2 in their vascular development, which might lead to an impairment of neuronal activity as a consequence since it is known that BBB breakdown is one of the fundamental biological events in Alzheimer’s disease^39^ and Huntington’s disease^53^ and other neurodegenerative and neurodevelopmental disorders^54,55^. Therefore, altered vascular impairment by MeCP2 mutation, which we showed in this paper, may presumably influence neurogenesis and brain microenvironment. Another literature showed neuronal activity also regulates blood-brain barrier efflux transport and cross-talk with each other ^56^. So far, there is little clear evidence to express the link between vascular impairment and regression of neurons in human Rett syndrome patients, although it was shown in mice model ^22^. This *in vitro* microvascular model from patient-derived cells and the capability of co-culture with neurons ^40^ could provide new tools to investigate the neurovascular interaction in the context of early developmental change in Rett syndrome for translational medicine.

In conclusion, we discovered miR-126-3p-mediated vascular alternation by MeCP2[R306C] mutation in Rett syndrome, along with downregulation of TEK and EDN1. The combination of our directed differentiation method into endothelial cells from patient-derived iPS cells with CRISPR/Cas knock in dox-inducible ETV2 and the microfluidic device facilitates to obtain robust 3D microvascular networks and allow us to measure permeability, which can recaptulate vasculogenesis process in Rett syndrome. Restoration of miR-126-3p level effectly rescure the endotehlail barrier function in RTT-iVascular along with modulating Ang/Tie2 signaling pathway and tight junction stabilization. This promised platform using patient-derived iPS cells and CRISPR/Cas9-based ETV2 knock-in system with the microfluidic device could apply to any other neurodevelopmental and neurodevelopmental disorder such as Alzheimer’s disease and amyotrophic lateral sclerosis.

## MATERIALS AND METHODS

### iPS culture from Rett syndrome patient

Rett syndrome patient-derived hiPS cells which carry heterozygous MECP2[R306C] (female, 8Y, missense, 4–7% of RTT patients) and its isogenic control was obtained from the Coriell Institute (WIC05i-127-325(MT) and WIC04i-127-33(WT)). Isogenic control was established by site-directed genome editing by CRISPR/Cas9 to fix the mutant allele of MECP2, resulting in isogenic control cells only having wild-type alleles of MECP2 in this line. All hiPS cells were maintained on Matrigel-coated 6-well plates in mTeSR plus medium (STEMCELL Technologies) with 10 μM Y-23632 (Rock inhibiter, STEMCELL technologies) for the first day after passages and subcultured every 5–7 days using ReLeSR reagent (STEMCELL Technologies). The pluripotency of iPS cells were characterized by immunostaining with OCT3/4, Nanog, TRA-1-60, and SSEA4 (**Supplementary fig. S 1**) The cells were cultured in incubator under 5% CO_2_ and 37 [C.

### Genome editing

To produce genome edited iPS cells line which express ETV2 for endothelial differentiation and hNGN2 under rtTA promoter, gene constructs were knock-in into the safe harbor AAVS1 locus of Rettsyndrome patient derived iPS cells and isogenic control iPS cells. For endothelial differentiation, pUCM-AAVS1-TO-hNGN2 (Addgene #105840) was used as backbone plasmid and digested with PacI (NEB) and NotI (NEB), then hNGN2 were replaced with ETV2 by Gibson cloning with the donor constructs with 40 bp overlap sequence (gBlock, IDT) to get pUCM-AAVS1-TO-ETV2. iPS cells were cultured in 6 well plates coated with Matrigel in mTeSR plus medium and detached using TypLE express for 5 min. A total of 2×10^6^ human iPS cells were co-transfected with 8 µg of PX458-AAVS1 plasmid which express SpCas9 and 6 µg of pUCM-AAVS1-TO-ETV2 or pUCM-AAVS1-TO-NGN2 in 100 µL of Opti-MEM. Electroporation was performed using a BTX ECM830 electroporator (Poring plus: voltage:150, pulse length: 5 msec, pulse: 100 msec; number of pulses: 2. Transfer pulse: voltage: 20V, pulse length: 100 msec, pulse: 100 msec; number of pulses: 5. After electroporation, iPS cells were subsequently seeded onto matrigel-coated plate in mTeSR plus with CloneR2. 24 hours post-electroporation, transfected iPS cells were subjected to the puromycin selection at 0.25 μg/ml with 10 μM of Y-27632. Then, the culture medium was replaced mTeSR plus and allowed to recover for one week. Once the cells reached to 80% confluent, the cells were dissociated by treatment with TrypLE Express (ThermoFisher Scientific) for 5 min. Then, the cell suspension was filtered with 70 um cell strainers (BD Falcon) in mTeSR plus medium. mCherry positive cells were sorted by cell sorter (BD Melody) and re-plated as a single clone to Matrigel-coated plate in mTeSR plus with CloneR2 to get a single colony. Expanded single colonies were then collected for genotyping.

PX458-AAVS1 was a gift from Adam Karpf (Addgene plasmid #113194; http://n2t.net/addgene:113194; RRID:Addgene_113194) and pUCM-AAVS1-TO-hNGN2 was a gift from Michael Ward (Addgene plasmid # 105840; http://n2t.net/addgene:105840; RRID:Addgene_105840).

### Genotyping

DNA were collected from genome edited iPS cells by QIAamp DNA kit (Qiagen). Gene fragment around AAVS1 locus were amplified by PCR to confirmed donor DNA sequence was correctly inserted to AAVS1 locus (**Supplementary** Fig 1.).

### Differentiation to endothelial cells

Genome edited iPS cells (ETV2-iPS cells) were plated onto Matrigel-coated 6 well-plate and cultured until the cells reached to 70-80% confluent. At day 0, the cells were treated with 1 ml of Accutase and incubated for 4 min, then add 1 mL of mTeSR plus for neuralization. Collected cells were further dissociated by pipetting up-and-down at 10 times with 1 mL of pipette tip to get single cells. Then, the cells were re-plated to Matrigel-coated 6 well-plate at 250k-350k cells/well in mTeSR plus medium and 10 μM Y-23632. Next day, the culture medium was replaced to mesoderm differentiation medium (DMEM/F12 supplemented with 1% GlutaMAX, 60 µg/ml of L-ascorbic acid, 6 µM of CHIR99021(GSK3 Inhibitor/WNT Activator, STEMCELL Technologies)) and cultured for 2 days with dairy medium change. Then, the culture medium was replaced with endothelial differentiation medium (DMEM/F12 supplemented with 1% GlutaMAX, 60 µg/mL of L-ascorbic acid, 50 ng/mL of VEGF-A, 50 ng/mL of bFGF2, 10ng/mL of EGF, and 10 uM of SB431542 (Activin/BMP/TGF-beta inhibitor, STEMCELL Technologies)), and 3 µg/mL of Doxycycline. Next day, the medium was changed to the same endothelial differentiation medium but without Doxycycline. At day 5, differentiated endothelial cells (iEC) were seeded to T150 culture flask for expansion culture in Vasculife VEGF Endothelial Medium Complete Kit (Lifeline cell technology, LL-0003) supplemented with 10% Fetal Bovine Serum (FBS), and 10 µM of SB431542, and then characterized flowcytometry.

### Differentiation to neurons

Genome edited iPS cells (NGN2-iPS cells) were plated onto Matrigel coated 6 well plate and cultured until the cells reached to 70-80% confluent. Then at day 0, the cells were treated with 1 ml of TrypLE Express and incubated for 5 min, then add 5 ml of mTeSR plus for neuralization. Collected cells were further dissociated by pipetting up-and-down at 10 times with 1ml of pipette tip to get a single cells. Then, the cells were re-plated to Matrigel-coated 6 well-plate at 200k cells/well in mTeSR plus medium and 10 μM Y-23632. Two days later, the culture medium was replaced to neuronal initiation medium differentiation medium (knockout DMEM/F12 supplemented with 15% KnockOut Serum replacement, 1% GlutaMAX, 1% NEAA, 10 uM of SB431542, 100 nM of LDN-193189 (TGF-beta/Smad inhibitor, STEMCELL Technologies), and 100 ng/ml of Doxycycline and cultured for 7 days with dairy medium change. Then, the cells were treated with 1 ml of Accutase and incubated for 10 min. Dissociated cells were spin down and resuspended in neuronal maturation medium (Neurobasal plus medium supplemented with 2% B27-plus, 1% of N2, and 100 ng/ml of Doxycycline (only initial 3 days) and seed to 6 of Matrigel-coated 6 well plate and cultured the cells for 10 days. Differentiated neurons were characterized by RT-PCR and used for Ca^2+^ imaging.

### Imaging-enhanced flowcytometry

To characterize differentiated iEC (RTT-iEC and WT-iEC) deeply with respect to the heterogeneity and tight junction expression, imaging-enhanced flow cytometry (Attune CytPix, Thermo Fisher Scientific) was performed. iECs were dissociated with TrypLE Express and fixed with 4% paraformaldehyde with 0.2% Triton X-100 and then the cells were filtered with 70 μm cell strainers (BD Falcon, 38030) in 1% BSA-PBS. Then, fixed cells were stained with Alexa Fluor 488 conjugated ZO-1 antibody (Thermo Fisher Scientific, ZO1-1A12, 1:100) and Hoechst 33342 for DNA staining for cells cycle analysis. In CytPix, the cells cell population were gated by FSC, SSC, and acquired images to get single cells. From acquired images, size of cells, perimeter, psudodiameter, entropy, and circularity of cells were computed in CytPix and integrated data (images, FSC, SSC, ZO-1 fluorescent value) were further analyzed in MATLAB (Mathworks).

### Engineering of 3D vascular networks

Both differentiated endothelial cells (iEC) at 8 ×10^6^ cells/ml and human lung fibroblasts (Lonza, CC-2512) at 1×10^6^ cells/ml as supporting cells were suspended in VascuLife containing thrombin (4 U/mL, Sigma). Cell suspension was further mixed with fibrinogen (3 mg/mL at final concentration, Sigma) on ice. The mixture was then introduced to the microfluidic device (idenTx 3, AIM Biotech) through the center channel. After introducing the gel mixture with cells, the device was incubated in incubator for 15 min to complete polymerization. Subsequently, VascuLife media supplemented with 10% Fetal Bovine Serum (FBS), and 10 µM of SB431542 were introduced into side medium channel in the devices. The cells were further cultured to allow microvasculature formation and the permeability assay was performed at day 7 from cell seeding.

### Permeability measurement in 3D vasculature and analysis

Vascular permeability was measured for various perfusable MVNs engineered in this study following published protocols ^39,41^. Briefly, 70 kDa Texas red dextran (Invitrogen) was perfused into the microvascular networks by generating a slight pressure gradient across the gel of the device. In detail, the culture medium in one media channel was first aspirated, followed by the injection of 10 µL of 10 µg/mL Texas red dextran solution. The process was then repeated for the other media channel, before imaging under a confocal microscope. Confocal images were captured with a 5 µm step size at 0 and 10 min, from which permeability was calculated. The detailed permeability measurement protocol and ImageJ Marco for permeability analysis can be found in previous protocol ^42^. Several regions of interest (ROIs) were captured via time-lapse volumetric imaging for each device.

### Quantification of mRNA and microRNA by real-time qRT-PCR

Total RNA was isolated from the cells from 2D culture using RNAeasy mini kit (Qiagen) following manufacture protocol. Total RNA was converted to cDNA using a Superscript VILO cDNA syntheses Kit (Thermofisher Scientific). qRT-PCR was performed with Quantstudio 3 (Applied biosystems) using Takara TB Green Premix Ex Taq II (Takara, SYBR Green). The primer sequences are shown in **Table S2.** The mRNA level of glyceraldehyde 3-phosphate dehydrogenase (GAPDH) was used as the internal standard in all experiments.

microRNA was isolated using miRNAeasy mini kit (Qiagen) and further enriched by excluding mRNA and tRNA. microRNA was modified for 3’ poly(A) tailing and 5’ ligation of an aptamer sequence by TaqMan Advanced miRNA cDNA Synthesis Kit (Thermo Fisher Scientific) to extend mature microRNAs present in the samples on each end prior to reverse transcription, and converted to cDNA. qRT-PCR was performed using TaqMan Advanced miRNA Assary (Assay ID: 477887_mir, for has-miR-126) and TaqMan Advanced miRNA human endogenous control (Thermo Fisher Scientific) with Quantstudio 3 (Applied biosystems). microRNA levels were normalized by as the combination of endogenous control as internal standard. The RT-PCR experiments were repeated at least three times with cDNAs prepared from separate cell culture along with experimental duplicates. Each reaction was assessed according to Melting curve to exclude abnormal amplification. Differential gene expressions were quantified by delta-delta Ct method. miR-126-3p binding domain were simulated by miRDB (https://mirdb.org/) ^34^and confirmed by TargetScanHuman ^58^ (https://www.targetscan.org/vert_80/). GC % and PhyloP score were obtained from UCSC Genome Browser (https://genome.ucsc.edu/).

### Characterization of neuronal activity by Ca^2+^ imaging

To characterize the direct impact of paracrine signals from iECs to neurons, we performed Ca2+ imaging using a two-photon microscope. At 3 weeks of culture, iPSC-derived NGN2-induced differentiated neurons were infected with rAAV-hSyn-jRGECO1a-WPRE and then cultured for at least 1 week after the infection. Twenty-four hours before the imaging, the culture medium was replaced with BrainPhys imaging optimization medium supplemented with 1% GlutaMAX, 2% B27 plus, 1% NEAA, and 1% Penicillin/Streptomycin. Time-lapse images were acquired using a two-photon microscope (Bruker) with a galvo-resonant scanner and an Insight X3 laser (Spectra-physics, 80 MHz) at 1040 nm for jRGECO1a excitation and a MaiTai laser (Spectra-physics, 80 MHz) at 860 nm for EGFP excitation. Both lasers were aligned with the same light path and were scanned at a sampling frequency of 60 Hz with a resolution of 512 x 512. The emission light was initially separated by a primary dichroic short pass filter mirror (890 nm), and then the green signal and red signal were separated by a dichroic mirror (Semrock, 540/50 nm) and further filtered by band pass filters (525/25 nm) and (610/45 nm), respectively. Both signals were detected using GaAsP photomultiplier tubes (H7422A-40, Hamamatsu, Japan). The microscope and laser were controlled by the Prairie View software (Bruker). Time-lapse images were first analyzed using the Suite2p package (Suite2p, python==3.10) to identify the signal neurons and segment them by determining regions of interest (ROIs). For each ROI time series, the baseline fluorescence was defined as the average of the lowest 10% of samples. ΔF/F was computed as (F - Faverage) / Faverage, and photobleaching was also normalized by the slope calculated by (Fend - Fstart) / time. The exported data (delta F) were loaded into MATLAB and further computed according to the analysis. To calculate spike frequency, delta F was initially deconvoluted to specify single neuron activity, and then spikes were detected using the “findpeaks” function in MATLAB. The spike frequency was calculated over a certain unit of time (minute). Correlation matrices were obtained by pairwise correlation among the neurons.

### Immunostaining and microarray

In 2D monolayer culture, the cells on the glass bottom dish were fixed with 4% paraformaldehyde in Phosphate-Buffer Saline (PBS) for 20 min at room temperature. Then the cells were permeabilized in 0.2% TritonX-100 in PBS for 5 min at room temperature and blocked in 1% BSA in PBS at 4 [C overnight. Primary antibody was then incubated with ZO-1 antibody (ThermoFisher, ZO1-1A12, 1:1000) at 4 [C overnight. After the cells were washed with PBS, secondary antibody was incubated with Goat anti-Rabbit IgG (H+L), Alexa Fluor 488 (1:1000) at 4 [C overnight. The cells then were incubated with Rhodamine phalloidin (70 nM) and Hoechst 33342 (1:10000) solution at room temperature for 10 min followed by the 3-time washing with PBS. Fluorescent images were acquired using confocal microscope (Olympus, FV1200) with 20X UPlanSApo objectives (NA: 0.75).

For 3D culture in microfluidic device, PBS were introduced into medium channel of microfluidic device and replaced 3 times for washing and removing the medium from microvascular networks. After removing excess PBS, cells were fixed with 4% paraformaldehyde in PBS for 4 hours at room temperature on rocking shaker. Then, the cells were permeabilized in 0.2% TritonX-100 in PBS for 1 hour at room temperature and blocked in 1% BSA in PBS at 4 [C overnight. Primary antibody (anti ZO-1 antibody, ThermoFisher, ZO1-1A12, 1:1000; anti-collagen IV mouse antibody, Abcam, ab19808, 1:500) was introduced into the device and incubated at 4 [C overnight on rocking shaker. Primary antibodies were carefully removed by washing with PBS for 1 hour and it repeated at least 3 times. Then, secondary antibodies (Goat anti-Rabbit IgG (H+L), Alexa Fluor 555 (1:1000); Goat anti-Mouse IgG (H+L), Alexa Fluor 647 (1:1000) were incubated at 4 [C overnight on rocking shaker. The cells then were incubated with Hoechst 33342 (1:10000) solution at room temperature overnight followed by the 3-time gentle washing with PBS. Z-stack images were acquired using confocal microscope (Olympus, FV1200) with 10X or 20X SApo objectives.

To quantify paracrine signals related to MMPs and TIMPs, the culture mediums were collected from cultured microfluidic devices. Then, protein expression in the medium were semi-quantitated by G-Series Human MMP Array 1 with manufacturer protocol.

### Statistical analysis

The reported values are the means of a minimum of three independent experiments. Data are presented as the mean ± SD. For equal variances and normality distribution, a Student’s t test was performed. To compare groups at multiple conditions, statistical comparisons were performed using one-way analysis of variance (ANOVA), with post hoc pairwise comparisons carried out using the Tukey-Kramer method. Statistical tests were performed using GraphPad Prism 9 (GraphPad Software, San Diego, CA). *p* values <0.05 or p values <0.01 were considered significant in all cases.

## Data availability

Read-level data from RNAseq will be deposited at GEO association. The plasmids used in this study will be deposited to Addgene. Any additional information required to re-analyses the data reported in this paper is available from the corresponding author upon request. The MATLAB for calculating pairwise correlation of neuronal activity and R code for RNAseq is available from GitHub.

## List of Supplementary Materials

Supplementary Fig. 1: Characterization of Rett syndrome patient-derived iPS cells.

Supplementary Fig. 2: RTT-iEC condition medium negatively impacted neuronal activity

Supplementary Fig. 3: Engieering BBB in vitro with iEC, pericyte and astrocyte

Supplementary Fig. 4: miR profiling and downregulated miR in RTT-iEC

Supplementary Table. 1: Key resources

Supplementary Table. 2: RT-PCR primer

## Author contributions

T.O, Z.W, R.D.K, and MS conceived and designed the experiments. TO, Z.W., and Y.J performed cell experiments and analyzed data. K.H. and M.C. performed cloning of EVT2 plasmid construct. TO, ZW, RDK, and MS RDK wrote the manuscript.

## Data and materials availability

Read-level data from RNA-seq have been deposited at GEO. Any additional information required to reanalyze the data reported in this paper is available from the corresponding author upon request.

## References and Notes

1 Lyst, M. J. & Bird, A. Rett syndrome: a complex disorder with simple roots. Nature Reviews Genetics 16, 261–275 (2015). 10.1038/nrg3897

2 Samaco, R. C. & Neul, J. L. Complexities of Rett Syndrome and MeCP2. The Journal of Neuroscience 31, 7951–7959 (2011). 10.1523/jneurosci.0169-11.2011

3 Chahrour, M. & Zoghbi, H. Y. The story of Rett syndrome: from clinic to neurobiology. Neuron 56, 422–437 (2007). 10.1016/j.neuron.2007.10.001

4 Sjöstedt, E. et al. An atlas of the protein-coding genes in the human, pig, and mouse brain. Science 367, eaay5947 (2020). doi:10.1126/science.aay5947

5 Karlsson, M. et al. A single–cell type transcriptomics map of human tissues. Science Advances 7, eabh2169 (2021). doi:10.1126/sciadv.abh2169

6 Lappalainen, R., Liewendahl, K., Sainio, K., Nikkinen, P. & Riikonen, R. S. Brain perfusion SPECT and EEG findings in Rett syndrome. Acta Neurologica Scandinavica 95, 44–50 (1997). 10.1111/j.1600-0404.1997.tb00067.x

7 Bianciardi, G. et al. Microvascular abnormalities in Rett syndrome. Clin Hemorheol Microcirc 54, 109–113 (2013). 10.3233/ch-131707

8 Saili, K. S. et al. Blood-brain barrier development: Systems modeling and predictive toxicology. Birth Defects Res 109, 1680–1710 (2017). 10.1002/bdr2.1180

9 Daneman, R., Zhou, L., Kebede, A. A. & Barres, B. A. Pericytes are required for blood–brain barrier integrity during embryogenesis. Nature 468, 562–566 (2010). 10.1038/nature09513

10 Ek, C. J., Dziegielewska, K. M., Habgood, M. D. & Saunders, N. R. Barriers in the developing brain and Neurotoxicology. Neurotoxicology 33, 586–604 (2012). 10.1016/j.neuro.2011.12.009

11 Saunders, N. R., Joakim Ek, C. & Dziegielewska, K. M. The neonatal blood-brain barrier is functionally effective, and immaturity does not explain differential targeting of AAV9. Nat Biotechnol 27, 804–805; author reply 805 (2009). 10.1038/nbt0909-804

12 Obermeier, B., Daneman, R. & Ransohoff, R. M. Development, maintenance and disruption of the blood-brain barrier. Nat Med 19, 1584–1596 (2013). 10.1038/nm.3407

13 Aschner, M. et al. Reference compounds for alternative test methods to indicate developmental neurotoxicity (DNT) potential of chemicals: example lists and criteria for their selection and use. Altex 34, 49–74 (2017). 10.14573/altex.1604201

14 Bal-Price, A. et al. Putative adverse outcome pathways relevant to neurotoxicity. Crit Rev Toxicol 45, 83–91 (2015). 10.3109/10408444.2014.981331

15 Corada, M. et al. The Wnt/beta-catenin pathway modulates vascular remodeling and specification by upregulating Dll4/Notch signaling. Dev Cell 18, 938–949 (2010). 10.1016/j.devcel.2010.05.006

16 Weis, S. M. & Cheresh, D. A. Pathophysiological consequences of VEGF-induced vascular permeability. Nature 437, 497–504 (2005). 10.1038/nature03987

17 Zlokovic, B. V. Neurovascular pathways to neurodegeneration in Alzheimer’s disease and other disorders. Nature Reviews Neuroscience 12, 723–738 (2011). 10.1038/nrn3114

18 Sweeney, M. D., Sagare, A. P. & Zlokovic, B. V. Blood–brain barrier breakdown in Alzheimer disease and other neurodegenerative disorders. Nature Reviews Neurology 14, 133–150 (2018). 10.1038/nrneurol.2017.188

19 Gray, M. T. & Woulfe, J. M. Striatal Blood–Brain Barrier Permeability in Parkinson’S Disease. Journal of Cerebral Blood Flow & Metabolism 35, 747–750 (2015). 10.1038/jcbfm.2015.32

20 Shaltiel-Karyo, R. et al. A Blood-Brain Barrier (BBB) Disrupter Is Also a Potent &#x3b1;-Synuclein (&#x3b1;-syn) Aggregation Inhibitor: A NOVEL DUAL MECHANISM OF MANNITOL FOR THE TREATMENT OF PARKINSON DISEASE (PD) *. Journal of Biological Chemistry 288, 17579–17588 (2013). 10.1074/jbc.M112.434787

21 Lee, R. L. & Funk, K. E. Imaging blood-brain barrier disruption in neuroinflammation and Alzheimer’s disease. Front Aging Neurosci 15, 1144036 (2023). 10.3389/fnagi.2023.1144036

22 Pepe, G. et al. Blood–Brain Barrier Integrity Is Perturbed in a Mecp2-Null Mouse Model of Rett Syndrome. Biomolecules 13, 606 (2023).

23 Panighini, A. et al. Vascular Dysfunction in a Mouse Model of Rett Syndrome and Effects of Curcumin Treatment. PLOS ONE 8, e64863 (2013). 10.1371/journal.pone.0064863

24 Pacheco, N. L. et al. RNA sequencing and proteomics approaches reveal novel deficits in the cortex of Mecp2-deficient mice, a model for Rett syndrome. Molecular Autism 8, 56 (2017). 10.1186/s13229-017-0174-4

25 Lyst, M. J. et al. Rett syndrome mutations abolish the interaction of MeCP2 with the NCoR/SMRT co-repressor. Nat Neurosci 16, 898–902 (2013). 10.1038/nn.3434

26 Luo, A. C. et al. A streamlined method to generate endothelial cells from human pluripotent stem cells via transient doxycycline-inducible ETV2 activation. Angiogenesis (2024). 10.1007/s10456-024-09937-5

27 Palikuqi, B. et al. Adaptable haemodynamic endothelial cells for organogenesis and tumorigenesis. Nature 585, 426–432 (2020). 10.1038/s41586-020-2712-z

28 Cui, N., Hu, M. & Khalil, R. A. Biochemical and Biological Attributes of Matrix Metalloproteinases. Prog Mol Biol Transl Sci 147, 1–73 (2017). 10.1016/bs.pmbts.2017.02.005

29 Issler, O. & Chen, A. Determining the role of microRNAs in psychiatric disorders. Nature Reviews Neuroscience 16, 201–212 (2015). 10.1038/nrn3879

30 Szulwach, K. E. et al. Cross talk between microRNA and epigenetic regulation in adult neurogenesis. Journal of Cell Biology 189, 127–141 (2010). 10.1083/jcb.200908151

31 Urdinguio, R. G. et al. Disrupted microRNA expression caused by Mecp2 loss in a mouse model of Rett syndrome. Epigenetics 5, 656–663 (2010). 10.4161/epi.5.7.13055

32 Tsujimura, K. et al. miR-199a Links MeCP2 with mTOR Signaling and Its Dysregulation Leads to Rett Syndrome Phenotypes. Cell Reports 12, 1887–1901 (2015). 10.1016/j.celrep.2015.08.028

33 van Solingen, C. et al. Antagomir-mediated silencing of endothelial cell specific microRNA-126 impairs ischemia-induced angiogenesis. J Cell Mol Med 13, 1577–1585 (2009). 10.1111/j.1582-4934.2008.00613.x

34 Liu, W. & Wang, X. Prediction of functional microRNA targets by integrative modeling of microRNA binding and target expression data. Genome Biology 20, 18 (2019). 10.1186/s13059-019-1629-z

35 Saharinen, P., Eklund, L. & Alitalo, K. Therapeutic targeting of the angiopoietin–TIE pathway. Nature Reviews Drug Discovery 16, 635–661 (2017). 10.1038/nrd.2016.278

36 Augustin, H. G., Young Koh, G., Thurston, G. & Alitalo, K. Control of vascular morphogenesis and homeostasis through the angiopoietin–Tie system. Nature Reviews Molecular Cell Biology 10, 165–177 (2009). 10.1038/nrm2639

37 Zhang, H., Yamaguchi, T., Kokubu, Y. & Kawabata, K. Transient ETV2 Expression Promotes the Generation of Mature Endothelial Cells from Human Pluripotent Stem Cells. Biological and Pharmaceutical Bulletin 45, 483–490 (2022). 10.1248/bpb.b21-00929

38 Wang, K. et al. Robust differentiation of human pluripotent stem cells into endothelial cells via temporal modulation of ETV2 with modified mRNA. Science Advances 6, eaba7606 10.1126/sciadv.aba7606

39 Campisi, M. et al. 3D self-organized microvascular model of the human blood-brain barrier with endothelial cells, pericytes and astrocytes. Biomaterials 180, 117–129 (2018). 10.1016/j.biomaterials.2018.07.014

40 Osaki, T., Sivathanu, V. & Kamm, R. D. Engineered 3D vascular and neuronal networks in a microfluidic platform. Scientific Reports 8, 5168 (2018). 10.1038/s41598-018-23512-1

41 Offeddu, G. S. et al. An on-chip model of protein paracellular and transcellular permeability in the microcirculation. Biomaterials 212, 115–125 (2019). 10.1016/j.biomaterials.2019.05.022

42 Hajal, C. et al. Engineered human blood–brain barrier microfluidic model for vascular permeability analyses. Nature protocols 17, 95–128 (2022). 10.1038/s41596-021-00635-w

43 Van den Steen, P. E., et al. Biochemistry and molecular biology of gelatinase B or matrix metalloproteinase-9 (MMP-9). Crit Rev Biochem Mol Biol 37, 375–536 (2002). 10.1080/10409230290771546

44 Visse, R. & Nagase, H. Matrix Metalloproteinases and Tissue Inhibitors of Metalloproteinases. Circulation Research 92, 827–839 (2003). 10.1161/01.RES.0000070112.80711.3D

45 Fanjul-Fernández, M., Folgueras, A. R., Cabrera, S. & López-Otín, C. Matrix metalloproteinases: Evolution, gene regulation and functional analysis in mouse models. Biochimica et Biophysica Acta (BBA) - Molecular Cell Research 1803, 3–19 (2010). 10.1016/j.bbamcr.2009.07.004

46 Wang, S. et al. The Endothelial-Specific MicroRNA miR-126 Governs Vascular Integrity and Angiogenesis. Developmental Cell 15, 261–271 (2008). 10.1016/j.devcel.2008.07.002

47 Wang, L. et al. Sensing and guiding cell-state transitions by using genetically encoded endoribonuclease-mediated microRNA sensors. Nature Biomedical Engineering (2024). 10.1038/s41551-024-01229-z

48 Qu, M.-J. et al. MicroRNA-126 is a prospective target for vascular disease. Neuroimmunology and Neuroinflammation 5, 10 (2018). 10.20517/2347-8659.2018.01

49 Lee, Y. et al. MicroRNA genes are transcribed by RNA polymerase II. Embo j 23, 4051–4060 (2004). 10.1038/sj.emboj.7600385

50 Liu, Y. et al. MECP2 directly interacts with RNA polymerase II to modulate transcription in human neurons. Neuron (2024). 10.1016/j.neuron.2024.04.007

51 Cheng, T.-L. et al. MeCP2 Suppresses Nuclear MicroRNA Processing and Dendritic Growth by Regulating the DGCR8/Drosha Complex. Developmental Cell 28, 547–560 (2014). 10.1016/j.devcel.2014.01.032

52 Nakashima, H. et al. MeCP2 controls neural stem cell fate specification through miR-199a-mediated inhibition of BMP-Smad signaling. Cell Reports 35, 109124 (2021). 10.1016/j.celrep.2021.109124

53 Di Pardo, A. et al. Impairment of blood-brain barrier is an early event in R6/2 mouse model of Huntington Disease. Sci Rep 7, 41316 (2017). 10.1038/srep41316

54 Barisano, G. et al. Blood–brain barrier link to human cognitive impairment and Alzheimer’s disease. Nature Cardiovascular Research 1, 108–115 (2022). 10.1038/s44161-021-00014-4

55 Aragón-González, A., Shaw, P. J. & Ferraiuolo, L. Blood-Brain Barrier Disruption and Its Involvement in Neurodevelopmental and Neurodegenerative Disorders. Int J Mol Sci 23 (2022). 10.3390/ijms232315271

56 Pulido, R. S. et al. Neuronal Activity Regulates Blood-Brain Barrier Efflux Transport through Endothelial Circadian Genes. Neuron 108, 937–952.e937 (2020). 10.1016/j.neuron.2020.09.002

57 Gan, Y. et al. Gut microbes in central nervous system development and related disorders. Front Immunol 14, 1288256 (2023). 10.3389/fimmu.2023.1288256

58 Agarwal, V., Bell, G. W., Nam, J.-W. & Bartel, D. P. Predicting effective microRNA target sites in mammalian mRNAs. eLife 4, e05005 (2015). 10.7554/eLife.05005

