## Supplemental file for "miR126-mediated impaired vascular integrity in Rett syndrome"

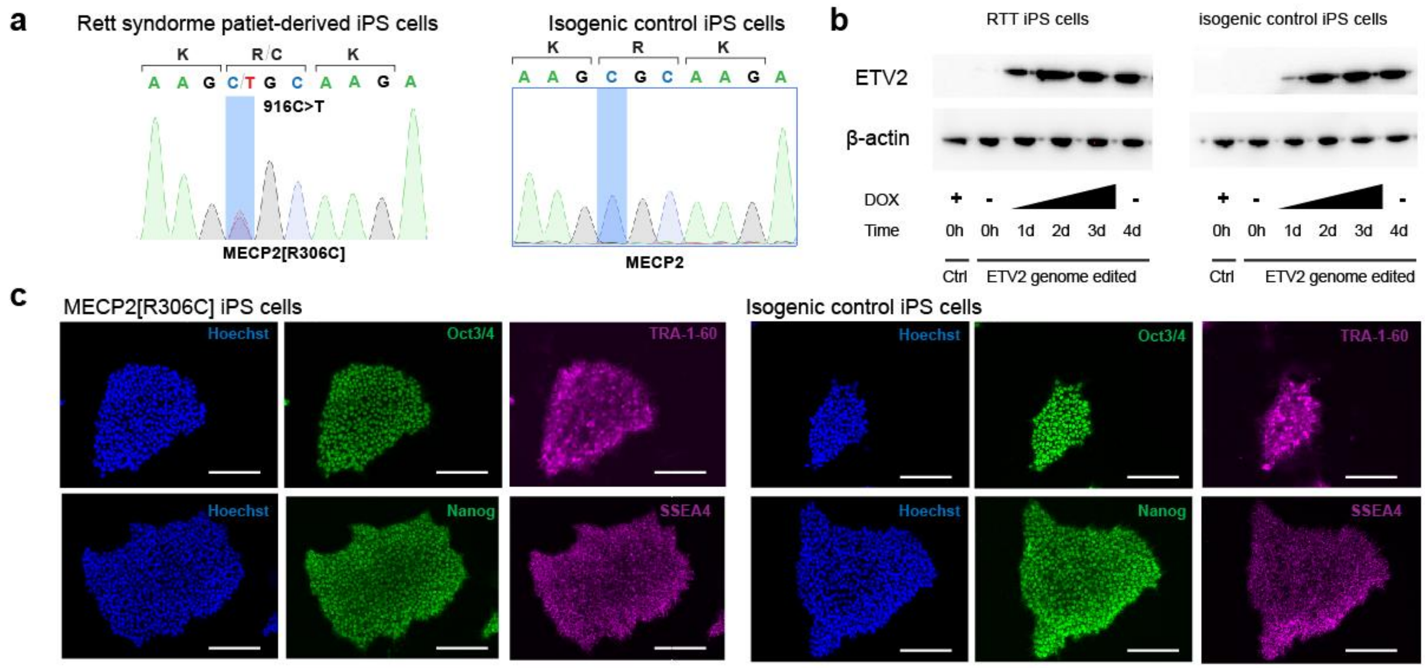


**Supplemental figure. 1 Characterization of Rett syndrome patient-derived iPS cells.** (**a**) Genotype to confirm MECP2 916c>T mutation in MeCP2[R306C] and isogenic control (**b**) ETV2 expression in the presence of DOX. (**c**) Immunostaining of OCT3/4, TRA-1-60 to show the pluripotency in iPS cells.

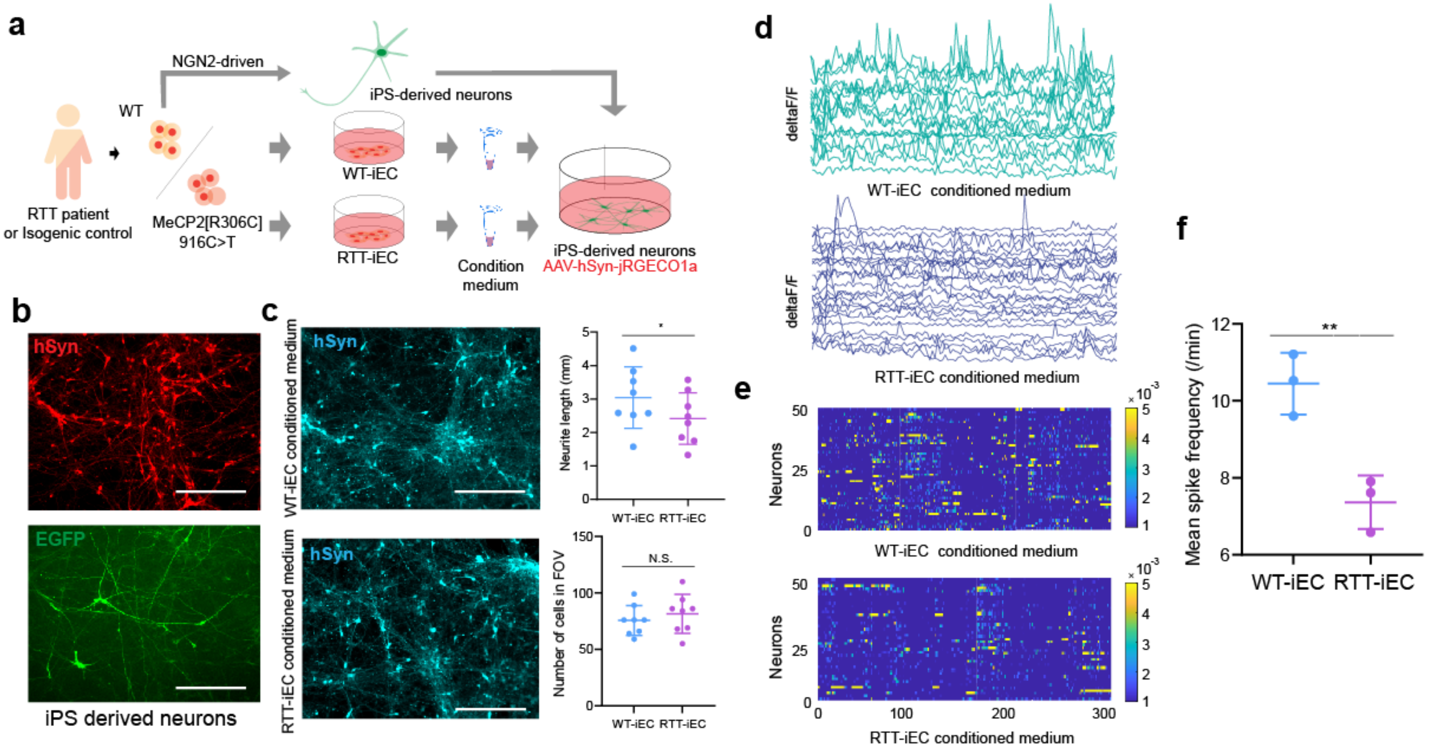


**Supplemental Fig. 2 RTT-iEC condition medium negatively impacted neuronal activity.** (**a**) Schematic illustration of treatment of WT-iNeurons with condition media from WT-iEC or RTT-iEC. (**b**) Immunostaining of hSyn to confirm differentiation to neurons from WT-iPS cells which express dox dependent NGN2. n =6. (**c**) Condition medium treatment did not affect morphological change with respect to neurite length and number of cells. (**d**) Calcium imaging was performed with AAV-hSyn-jRGECO1a. (**e, f**) Condition medium from RTT-iEC down reulgated neuronal activity. n =3. Student's t-test, **, *p*<0.01, * *p*<0.05.

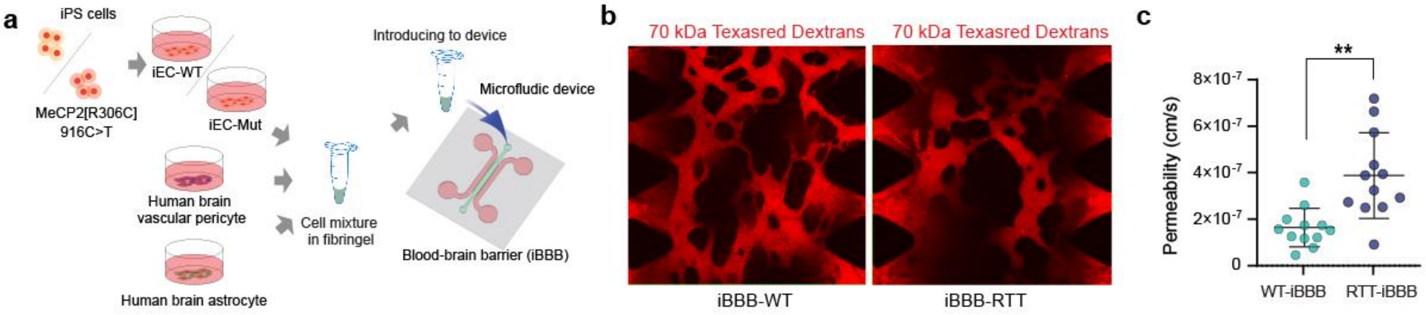


**Supplemental Fig. 3 Engieering BBB in vitro with iEC, pericyte and astrocyte.** (**a**) Schematic illustration of formation of BBB by co-cultureing, endothelial cells, pericytes, and astrocytes in fibrin gel. (**b**) BBB formation from Rett syndrome patient-derived iEC and isogenic control iEC. (**c**) RTT-iBBB displayed higher permeability compared to control BBB. n =12. Student's t-test, **, $p$<0.01, * $p$<0.05.

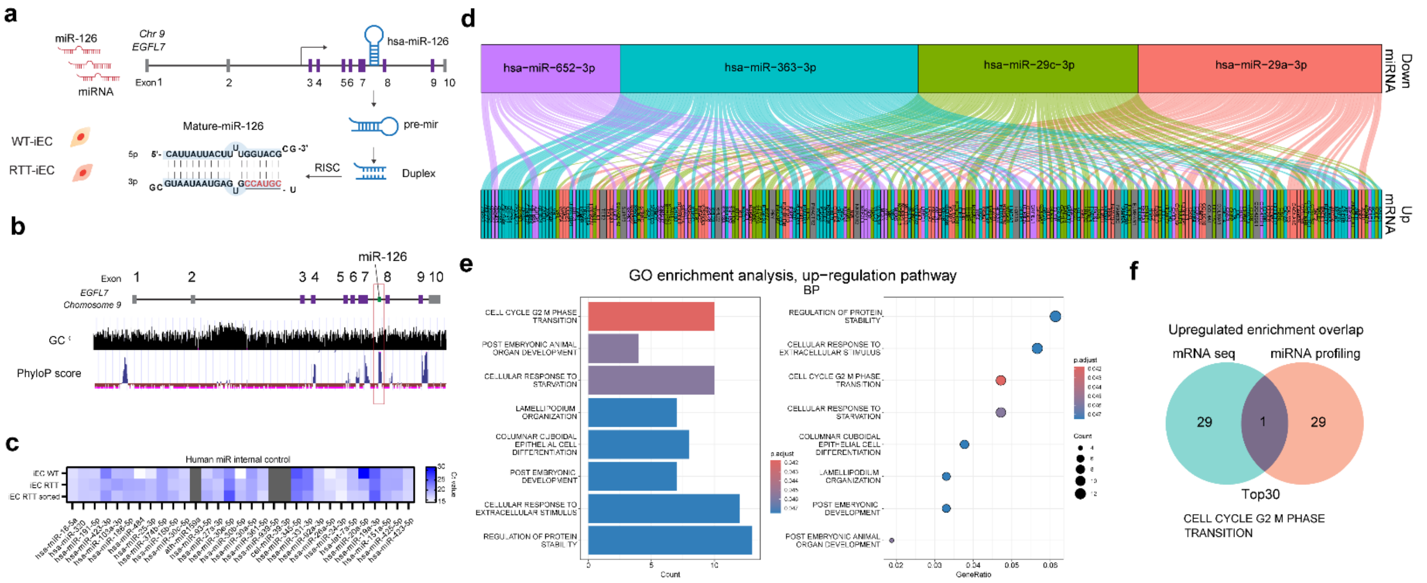

**Supplemental Fig. 4 miR profiling and downregulated miR in RTT-iEC.** (**a, b**) miR126-3p is endothelial specific microRNA, which is located in intron of EGFL7 and has higher PhyloP score. (**c**) microRNA internal control that used for normalization of target miR expression. (**d**) four down-regulated miR in RTT-iEC and target gene predicted by miRDB. (**e**) GO enrichment analysis by upregulated genes obtained from (**d**). (**f**) miRNA-mRNA integrated analysis revealed that overwrapped sinaling pathway (Cell cycle G2 M phase transition).

**Supplemental table S1. Key resources**

| Reagent and resource | Source | Identifier |
|---|---|---|
| **Cell line** | | |
| Rett syndrome patient derived iPS cells (female, 8Y, missence) | Coriell Institute | WIC05i-127-325(MT) |
| Isogenic control pair, wild type | Coriell Institute | WIC04i-127-33(WT) |
| AAVpro 293T | Takara | N.A. |
| **Plasmid** | | |
| pUCM-AAVS1-TO-hNGN2 | Addgene | # 105840 |
| pUCM-AAVS1-TO-hETV2 | This study | N.A. |
| PX458-AAVS1 | Addgene | #113194 |
| pRSV-Rev | Addgene | #12253 |
| pMDLg/pRRE | Addgene | #12251 |
| pMD2.G | Addgene | #12259 |
| miRZip-126-3p anti-miR-126-3p microRNA construct | SBI | MZIP126-3p-PA-1 |
| miRZip™ & pGreenPuro™ shRNA Scramble Hairpin Negative Control | SBI | MZIP000-PA-1 |

**Supplemental table S2.**

| Gene | Forward primer 5'-3' | Reverse primer 5'-3' |
|---|---|---|
| **CD31** | AACAGTGTTGACATGAAGAGCC | TGTAAAACAGCACGTCATCCTT |
| **ZO-1** | CAACATACAGTGACGCTTCACA | CACTATTGACGTTTCCCCACTC |
| **OCLDN** | ACAAGCGGTTTTATCCAGAGTC | GTCATCCACAGGCGAAGTTAAT |
| **CLDN5** | CTCTGCTGGTTCGCCAACAT | CAGCTCGTACTTCTGCGACA |
| **PGP** | TGACCCGCACTTCAGCTAC | GGGCTTCCCGATGATGTCG |
| **LRP1** | AGCCAGCTATGCACCAACAC | CCTTGCAGGAGCGGTTATC |
| **LAT1** | CCGTGAACTGCTACAGCGT | CTTCCCGATCTGGACGAAGC |
| **hTfR** | GGCTACTTGGGCTATTGTAAAGG | CAGTTTCTCCGACAACTTTCTCT |
| **GLUT1** | TCTGGCATCAACGCTGTCTTC | CGATACCGGAGCCAATGGT |
| **ABCA** | ACCCACCCTATGAACAACATGA | GAGTCGGGTAACGGAAACAGG |
| **CAT1** | ATCATCGGTACTTCAAGCGTAGC | GGCGTTCAGAGTCATGTGTGT |
| **MRP1** | AAGGAGGTACTAGGTGGGCTT | CCAGTAGGACCCTTCGAGC |
| **MARP4** | AGCTGAGAATGACGCACAGAA | ATATGGGCTGGATTACTTTGGC |
| **MCT1** | AGGTCCAGTTGGATACACCCC | GCATAAGAGAAGCCGATGGAAAT |
| **EGFL7** | TGCAGACGGTACACTCTGTGTG | TGCAGCCTCTGCACTTCTTCCT |
| **IGFBP2** | CGAGGGCACTTGTGAGAAGCG | TGTTCATGGTGCTGTCCACGTG |
| **VEGFA** | AGTCTGTCGTCCAGTAGCACCA | ACTGGAGCCATACTCATCCGAG |
| **NFKBIA** | TCCACTCCATCCTGAAGGCTAC | CAAGGACACCAAAAGCTCCACG |
| **HIF1a** | TATGAGCCAGAAGAACTTTTAGGC | CACCTCTTTTGGCAAGCATCCTG |
| **ADAM9** | CTTGCTGCGAAGGAAGTACCTG | CACTCACTGGTTTTTCCTCGGC |
| **PIK3R2** | CAGTACAACGCCAAGCTGGACA | TGCTGGTGGTAGACCTTGAGCT |
| **SPRED1** | CAGCCAGGCTTGGACATTCA | TGGGACTTTAGGCTTCCACAT |
| **TEK** | TTAGCCAGCTTAGTTCTCTGTGG | AGCATCAGATACAAGAGGTAGGG |
| **EDN1** | AGAGTGTGTCTACTTCTGCCA | CTTCCAAGTCCATACGGAACAA |
| **FOXO1** | TGATAACTGGAGTACATTTCGCC | CGGTCATAATGGGTGAGAGTCT |
| **ANGPT2** | CTCGAATACGATGACTCGGTG | TCATTAGCCACTGAGTGTTGTTT |
| **GAPDH** | TGT GGG CAT CAA TGG ATT TGG | ACA CCA TGT ATT CCG GGT CAA T |
